# Single-cell tumor phylogeny inference with copy-number constrained mutation losses

**DOI:** 10.1101/840355

**Authors:** Gryte Satas, Simone Zaccaria, Geoffrey Mon, Benjamin J. Raphael

## Abstract

**Motivation:** Single-cell DNA sequencing enables the measurement of somatic mutations in individual tumor cells, and provides data to reconstruct the evolutionary history of the tumor. Nearly all existing methods to construct phylogenetic trees from single-cell sequencing data use single-nucleotide variants (SNVs) as markers. However, most solid tumors contain copy-number aberrations (CNAs) which can overlap loci containing SNVs. Particularly problematic are CNAs that delete an SNV, thus returning the SNV locus to the unmutated state. Such mutation losses are allowed in some models of SNV evolution, but these models are generally too permissive, allowing mutation losses without evidence of a CNA overlapping the locus.

**Results:** We introduce a novel *loss-supported* evolutionary model, a generalization of the infinite sites and Dollo models, that constrains mutation losses to loci with evidence of a decrease in copy number. We design a new algorithm, <u>S</u>ingle-<u>C</u>ell <u>A</u>lgorithm for <u>R</u>econstructing the <u>L</u>oss-supported <u>E</u>volution of <u>T</u>umors (Scarlet), that infers phylogenies from single-cell tumor sequencing data using the loss-supported model and a probabilistic model of sequencing errors and allele dropout. On simulated data, we show that Scarlet outperforms current single-cell phylogeny methods, recovering more accurate trees and correcting errors in SNV data. On single-cell sequencing data from a metastatic colorectal cancer patient, Scarlet constructs a phylogeny that is both more consistent with the observed copy-number data and also reveals a simpler monooclonal seeding of the metastasis, contrasting with published reports of polyclonal seeding in this patient. Scarlet substantially improves single-cell phylogeny inference in tumors with CNAs, yielding new insights into the analysis of tumor evolution.

**Availability:** Software is available at github.com/raphael-group/scarlet

## 1 Introduction

Cancer arises from an evolutionary process during which somatic mutations accumulate in a population of cells. Different cells within a tumor acquire distinct complements of somatic mutations, resulting in a heterogeneous tumor. Quantifying this intra-tumor heterogeneity and reconstructing the evolutionary history of a tumor is crucial for diagnosis and treatment of cancer^1, 2^. The evolution of a tumor is typically described by a *phylogenetic tree*, or *phylogeny*, whose leaves represent the cells observed at the present time and whose internal nodes represent ancestral cells. Tumor phylogenies are challenging to reconstruct using DNA sequencing data from bulk tumor samples, since this data contains mixtures of mutations from thousands–millions of heterogeneous cells in the sample^3–15^. Recently, single-cell DNA sequencing (scDNA-seq) of tumors has become more common, and new technologies such as those from 10X Genomics^16^, Mission Bio^17^, and others^18–20^ are improving the efficiency and lowering the costs of isolating, labeling, and sequencing individual cells. While scDNA-seq overcomes the difficulties of phylogeny reconstruction from bulk samples, it introduces a new challenge of higher rates of missing data and errors due to DNA amplification errors, undersampling, and sequencing errors^18^.

Early work in phylogeny inference from scDNA-seq data uses single-nucleotide variants (SNVs) as phylogenetic markers. A particular challenge for SNV-based analysis is high rates (up to 30% for high-depth scDNA-seq^18^) of *allele dropout* errors, where only one of two alleles is observed at a heterozygous site. Methods address this challenge by using an evolutionary model to infer a phylogeny while simultaneously imputing missing data and correcting errors in the observed SNVs. Algorithms such as SCITE^21^, On-coNEM^22^, SciΦ^23^, and B-SCITE^24^ use the simplest phylogenetic model for SNVs, the *infinite sites model*. In this model, a locus in a cell has one of two states: an SNV (or mutation) iseither present at the locus (state 1) or absent (state 0). Transitions between states are constrained in the phylogeny such that each mutation is gained (0 → 1) at most once during evolution, and never subsequently lost (1 → 0). A phylogeny that respects the infinite sites model is known as a *perfect phylogeny* and the state of mutations in the leaves of the phylogeny is summarized by a *mutation matrix* whose binary entries indicate the presence (state 1) or absence (state 0) of every mutation in each observed cell (Fig. 1(A)). On error-free data, the perfect phylogeny is unique^25^. However, on typical scDNA-seq data, errors in the mutation matrix must be corrected to yield a perfect phylogeny model. Because many such corrections are possible, multiple phylogenies are typically equally consistent with the data (Fig. 1(B)).

**Fig. 1:**
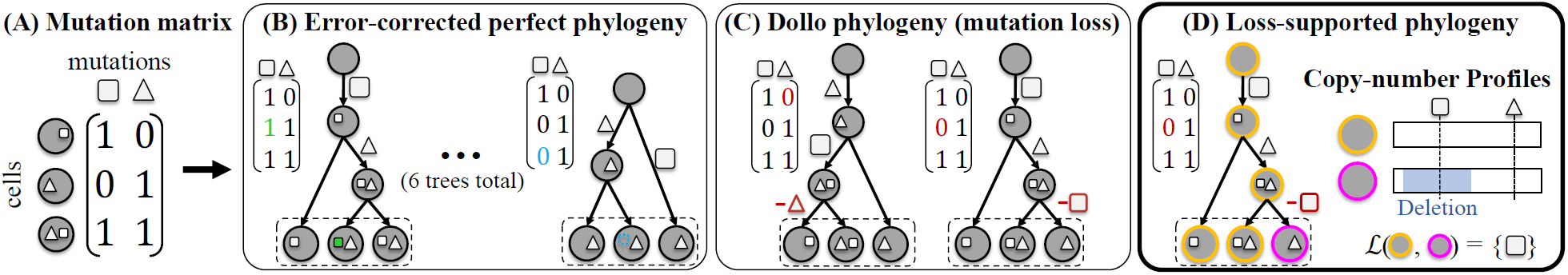
Loss-supported model reduces ambiguity in inference of phylogeny from single-cell sequencing data. (A) A mutation matrix with two mutations in three cells does not admit a perfect phylogeny. This may be due to either errors or mutation losses. (B) Under the infinite sites model, existing methods correct errors in the observed matrix to yield a perfect phylogeny. (C) Under the Dollo model, existing methods identify mutation losses to explain violations of the infinite sites model. Both the infinite sites and Dollo models yield multiple equally-plausible phylogenies. (D) The loss-supported model overcomes this ambiguity by using copy-number data to constrain mutation losses.

An additional challenge in inferring phylogenies from cancer sequencing data is that somatic mutations in tumors occur across all genomic scales from SNVs to copy-number aberrations (CNAs), which amplify or delete larger genomic regions. CNAs may overlap SNVs and affect the state of SNVs in cells; e.g., a deletion that overlaps an SNV may result in a *mutation loss* (1 → 0). The infinite sites model does not allow mutation losses and therefore may yield incorrect phylogenies when applied to SNVs in regions containing CNAs. One solution is to exclude regions containing CNAs and build phylogenies from SNVs in diploid, or copy-neutral, regions. However, ≈90% of solid tumors are highly aneuploid^26^, containing extensive CNAs, and ≈30% of solid tumors have whole-genome duplications^27^. Identifying collections of SNVs with no possibility of overlapping CNAs during evolution of such tumors may be challenging.

Recently, several methods^28–33^ have been introduced for single-cell phylogeny inference that allow loss of mutations. SPhyr^28^, SASC^29^, and PyDollo^30^ use the *Dollo model*^34^, which relaxes the infinite sites model. In the Dollo model, a mutation may be gained (0 → 1) at most once, but may be lost (1 → 0) multiple times. SiFit^31^, SiCloneFit^32^, and PhiSCS^33^ use the finite sites model, a further relaxation that allows mutation to be gained more than once. A challenge in using these less stringent evolutionary models is that they increase the ambiguity in phylogenetic reconstruction (Fig. 1(C)). Even in simple cases with no error, multiple phylogenies are consistent with the data and the number of phylogenies further increases when there are errors and uncertainty in the mutation matrix. Both the errors in scDNA-seq data and the mutation losses in the phylogeny conspire to yield considerable challenges and ambiguity in the single-cell phylogeny inference problem. This ambiguity is further amplified because both sequencing errors and losses result in the same signal in the observed data: an observed ‘0’ in the mutation matrix instead of a ‘1’. Thus, it is particularly difficult to distinguish between errors in the data and potential mutation losses.

A major limitation in using the Dollo or finite sites models to allow mutation losses is that neither of these models consider evidence from CNAs that support or refute a mutation loss at a locus. While more general multi-state models of tumor evolution have been used to infer phylogenies from bulk tumor sequencing data^7–9^, these approaches neither model the errors in scDNA-seq data nor scale to hundreds–thousands of observed cells. Since mutation losses are the major complication in SNV evolution and responsible for most of the violations of the infinite sites model in scDNA-seq data^30, 35^, the full generality of a multi-state model may not be necessary to obtain accurate phylogenies from scDNA-seq data. Rather, we describe an approach that constrains mutation losses by using copy-number data from the same cells.

We introduce Scarlet (<u>S</u>ingle-<u>C</u>ell <u>A</u>lgorithm for <u>R</u>econstructing the <u>L</u>oss-supported <u>E</u>volution of <u>T</u>umors), an algorithm that infers phylogenies from scDNA-seq data by integrating SNVs and copy-number data. Scarlet is based a novel evolutionary model, the *loss-supported phylogeny*, that constrains mutation losses to loci where the copy-number data has evidence of a deletion (Fig. 1(D)). The loss-supported phylogeny generalizes the infinite sites and Dollo models. Scarlet also relies on a probabilistic model of the read counts for each SNV to address errors and missing data that are common in scDNA-seq. On simulated data, we show that Scarlet infers more accurate phylogenies compared to existing methods. We then use Scarlet to analyze scDNA-seq data from a metastatic colorectal cancer patient from Leung et al. (2017)^36^. We find that the published phylogeny – constructed from SNVs under the infinite sites model – has the implausible conclusion that genome-wide copy-number profiles evolved twice independently during the evolution of this tumor. In contrast, Scarlet infers a loss-supported phylogeny that has three mutation losses, with each loss supported by a copy-number change at the locus. Moreover, the Scarlet phylogeny supports the hypothesis of a single migration between the colon primary tumor and liver metastasis (*monoclonal seeding*). In contrast, previous published phylogenies^32, 36^ reported a more complex origin of the metastasis with multiple migrations (*polyclonal seeding*). By integrating information from both SNVs and CNAs, Scarlet obtains more accurate reconstructions of tumor evolution at single-cell resolution.

## 2 Results

### 2.1 SCARLET algorithm for Loss-supported Phylogeny Model

We developed a new algorithm, Scarlet (<u>S</u>ingle-<u>C</u>ell <u>A</u>lgorithm for <u>R</u>econstructing the <u>L</u>oss-supported <u>E</u>volution of <u>T</u>umors) to infer phylogenetic trees from single-cell DNA sequencing (scDNA-seq) data by integrating data from both single-nucleotide variants (SNVs) and copy-number aberrations (CNAs). Scarlet has three important features (Fig. 2): (1) a novel evolutionary model, the *loss-supported phylogeny*, which constrains mutation losses to loci where there is a corresponding decrease in copy number; (2) an algorithm to compute a loss-supported phylogeny by *refinement* of a coarse phylogenetic tree derived from copy-number data alone; (3) maximum-likelihood inference of SNVs using a probabilistic model of observed read counts in scDNA-seq data. We describe each of these key features below.

**Fig. 2:**
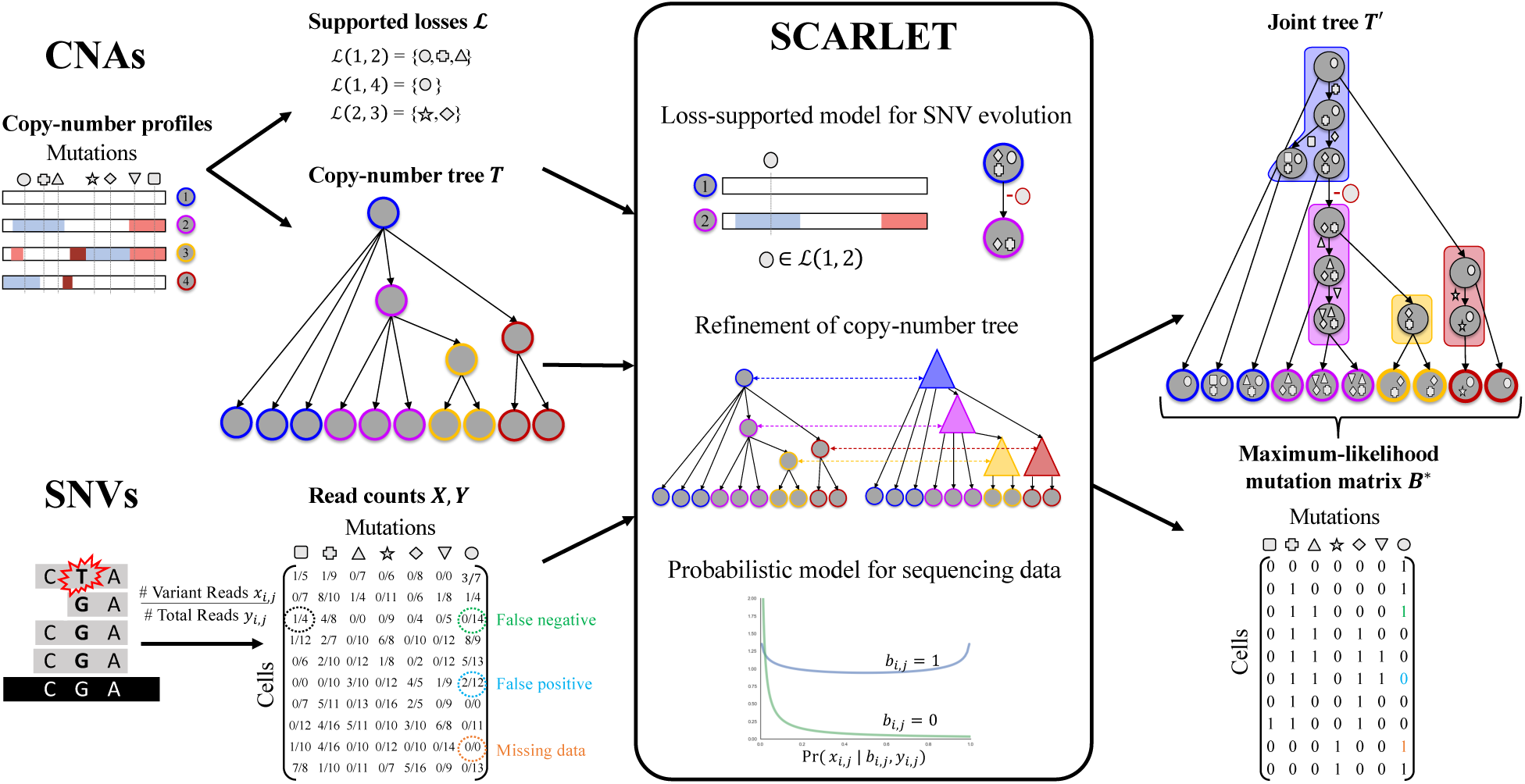
Scarlet algorithm for the Maximum Likelihood Loss-Supported Refinement problem. Scarlet integrates single-nucleotide variants (SNVs) and copy-number aberrations (CNAs) for tumor phylogeny inference. For CNAs, observed copy-number profiles indicate amplified (red) or deleted (blue) genomic regions along the entire genome and are used to obtain two inputs for Scarlet. First, supported loss sets ℒ (*c, c*′) for pairs of copy-number profiles (empty sets are not shown) indicate mutations that are affected by deletions. Second, a copy-number tree *T* which describes the ancestral relationships between observed cells (leaves) as determined by copy-number profiles. For SNVs, variant **X** and total **Y** read counts are provided to Scarlet for every cell and every mutation. Scarlet computes a joint tree *T*′ on the observed cells and a maximum-likelihood mutation matrix *B*\* by constraining mutation losses to the supported loss sets ℒ, computing a refinement *T*′ of *T*, and selecting the maximum-likelihood **B**\* using a probabilistic model for the presence (*b*_*i,j*_ = 1) or absence (*b*_*i,j*_ = 0) of each SNV in each cell.

The loss-supported model is a model of SNV evolution where mutation gains (0 → 1) occur at most once, but mutation losses (1 → 0) are constrained by sets ℒ of *supported losses* that are defined by CNAs in the same cells (Fig. 1(D)). Specifically, we assume that for each cell we measure both a mutation profile **b** of SNVs *and* a *copy-number profile c*. For each pair (*c, c*′) of copy-number profiles, we define the *supported loss set* ℒ (*c, c*′) as the set of SNVs at loci where there is a *decrease* in copy number (e.g., due to a deletion or loss-of-heterozygosity (LOH) event) between profiles *c* and *c*′. In the loss-supported phylogeny, a mutation loss at an SNV loci *a* is allowed between cells *v* and *w* only if *a* is in ℒ (*c*_*v*_, *c*_*w*_). The loss-supported model can thus be viewed as a generalization of other models for SNV evolution: the perfect phylogeny model is the special case where ℒ = ≈, while the Dollo model and finite sites model corresponds to ℒ being the complete set of all mutations. In contrast to these extremes, the loss-supported model allows for intermediate values of ℒ derived from copy-number data.

The loss-supported model depends on the copy-number profiles of both the observed and ancestral cells. However, we do not directly measure the copy-number profiles of the ancestral cells. To overcome this limitation, Scarlet uses a *copy-number tree T* which is derived from the copy-number profiles of the observed cells^13, 37–39^ (Fig. 2). Scarlet computes the supported loss sets ℒ from the copy-number profiles of the observed cells (leaves of *T*) and the copy-number profiles of the ancestral cells (internal vertices of *T*). Typically, scDNA-seq data of SNVs (e.g., from targeted sequencing) measures copy-number profiles with low resolution, and thus tumor cells share a limited number of distinct copy-number profiles. Consequently, the copy-number tree *T* has many multifurcations, or unresolved ancestral vertices with more than two children. Scarlet finds a joint tree *T*′ that is a loss-supported phylogeny and a *refinement*^40^*of T* by resolving multifurcations in *T* using the mutation profiles of the observed cells (Fig. 2).

Data from scDNA-seq typically has high error rates in identifying SNVs, and particularly high rates of false negatives and missing data due to amplification bias and allele dropout^18^. Scarlet models these errors using a beta-binominal distribution^23^ of the observed read counts. As such, Scarlet computes the loss-supported refinement *T*′ that maximizes the likelihood of the observed sequencing data under this probabilistic model (Fig. 2).

### 2.2 Simulated Data

We compared Scarlet to four existing algorithms that build phylogenies from single-cell sequencing data, SCITE^21^, SciΦ^23^, SPhyR^28^, and SiFit^31^, on simulated data. We simulated 50 trees, each with 20 mutations, 4 copy-number profiles, and 1-8 mutation losses per tree. From these trees, we simulated 100 observed cells with each cell having equal probability of being a child of any vertex in the simulated tree, and simulated sequencing data with an expected sequencing depth of 100× and allelic dropout rate of 0.15. Additional details of simulated data and parameters of each method are in Section S2.4.

We evaluated the phylogenies output by the methods by two measures that have been previously used in tumor evolution studies^11, 15, 23, 28, 29, 41^. First, the *mutation matrix error* 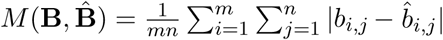 is the normalized Hamming distance between the inferred binary mutation matrix 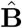 and the true binary mutation matrix **B** and assesses the accuracy of the error-corrected mutation profiles for each observed cell. Second, the *pairwise ancestral relationship error* 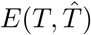 is the proportion of pairwise ancestral relationships between mutations in the inferred tree 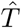 that differ from the ancestral relationships in the true tree *T*. Specifically, every pair *a, a*′ of mutations has one of four possible ancestral relationships in 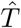 and in *T*: (1) *a* and *a*′ occur on the same branch; (2) *a* is ancestral to *a*′; (3) *a*′ is ancestral to *a*; (4) *a* and *a*′ are incomparable. Note that we do not calculate the tree error for SiFit because it uses a finite sites model which allows mutations to recur and consequently pairs of mutations may not have a unique relationship.

Scarlet outperforms all other methods on both mutation matrix error and ancestral relationship error (Fig. 3(A)-(B)). The high errors of SCITE and SciΦ were expected since these methods use an infinite sites model while the simulations include mutation losses which violates the model assumptions. However, the methods that do allow mutation losses, SPhyR (based on the *k*-Dollo model) and SiFit (based on the finite sites model), do not exhibit improvement over the other methods and perform worse than Scarlet. These results confirm that models that include unconstrained mutation losses have significant ambiguity as it is difficult to distinguish between true mutation losses and false positives/negatives in the data (Fig. 1). By using copy-number information to constrain mutation losses, Scarlet overcomes the ambiguity in phylogeny reconstruction and obtains lower error in the inferred mutation matrix and phylogeny.

**Fig. 3:**
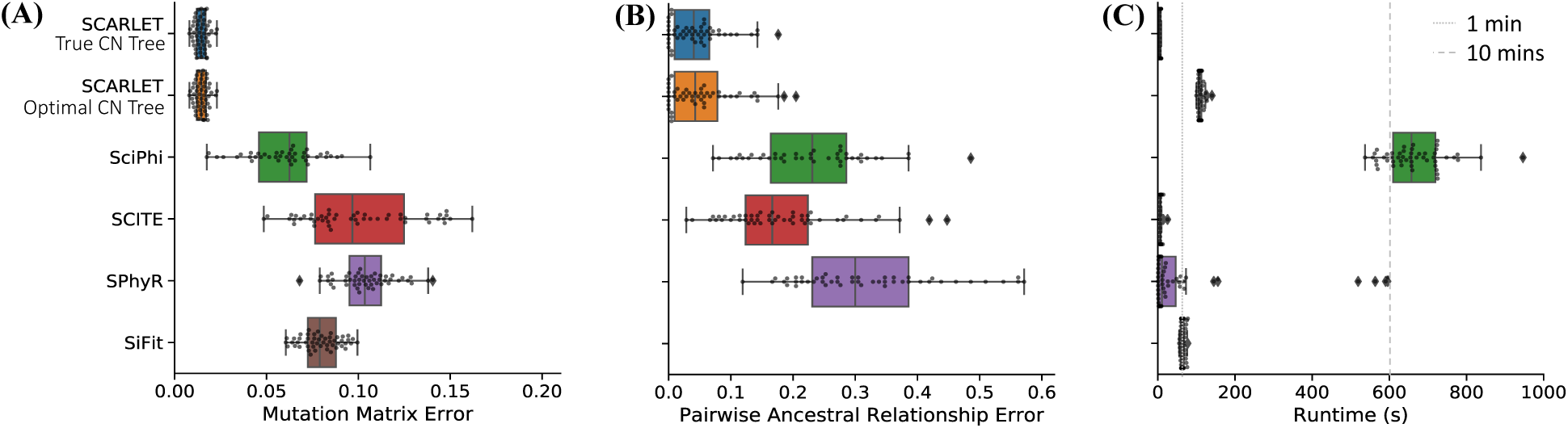
Scarlet outperforms existing methods for phylogeny inference on simulated single-cell data. (A) Mutation matrix error, (B) Pairwise ancestral relationship error, and (C) Runtime for each method. Scarlet was run either knowing (‘True CN Tree’) or not knowing (‘Optimal CN Tree’) the true copy number tree.

We evaluated the effect of the input copy number tree on Scarlet’s accuracy by running Scarlet in two modes: when the true copy-number tree is either known (‘Scarlet True CN Tree’) or unknown (‘Scarlet Optimal CN Tree’). In this latter case, we enumerated all copy-number trees, ran Scarlet once for each copy-number tree, and output the solution with the highest likelihood. In both cases, we provided Scarlet with the true copy-number profiles of each cell and the true set ℒ of supported losses. Scarlet exhibited comparable performance when running with or without knowledge of the copy-number tree (Fig. 3(A)-(B)). Notably, in 46/50 simulated instances, the maximum likelihood solution obtained when running Scarlet with unknown copy-number tree was identical to the solution found when providing the true copy-number tree. Clearly, running Scarlet with all possible copy-number trees (16 copy-number trees in this simulation) increases the runtime (Fig. 3(C)), but the runtime remains reasonable when the number of copy-number profiles is small, which is the case for many real datasets (see below).

### 2.3 Single-cell phylogeny of metastatic colorectal cancer

We used Scarlet to analyze single-cell DNA sequencing of a metastatic colorectal cancer patient CRC2 from Leung et al^36^. This data set included targeted sequencing of 1000 genes in 141 cells from a primary colon tumor and 45 cells from a matched liver metastasis (Fig. S1(A)). The authors identified 36 single-nucleotide variants (SNVs) and used SCITE^21^ to derive a perfect phylogeny from these SNVs (Fig. 4(A)). This perfect phylogeny tree shows two distinct branches of metastatic cells, and Leung et al.^36^ concluded that this was evidence of *polyclonal seeding* of the liver metastasis; i.e., two distinct cells (or groups of cells) with different complements of mutations migrated from the primary colon tumor to the liver metastasis. Examining the copy-number data, one finds a curious discrepancy between the SCITE tree and the single-cell copy-number profiles. Whole-genome sequencing of 42 single cells from the same patient reveals that all metastatic cells share losses of chromosomes 2, 3p, 4, 7, 9, 16, 22 relative to the cells in the primary tumor (Fig. 4(B)). According to the SCITE tree, all of these large copy-number aberrations (CNAs) would had to have occurred *twice independently* in the two distinct branches of metastatic cells. Although CNAs can exhibit homoplasy, this high rate of occurrence of the exact same events seems highly unlikely. Thus, we observe an inconsistency between the copy-number data and the SCITE tree constructed using only SNV data. Notably, this same dataset was recently analyzed by SiCloneFit^32^ using a finite sites model. The SiCloneFit tree also showed two branches of metastastic cells and concluded that there was polyclonal seeding of the metastases. Thus, the SiCloneFit phylogeny also has the same inconsistency between the SNV phylogeny and copy-number data.

**Fig. 4:**
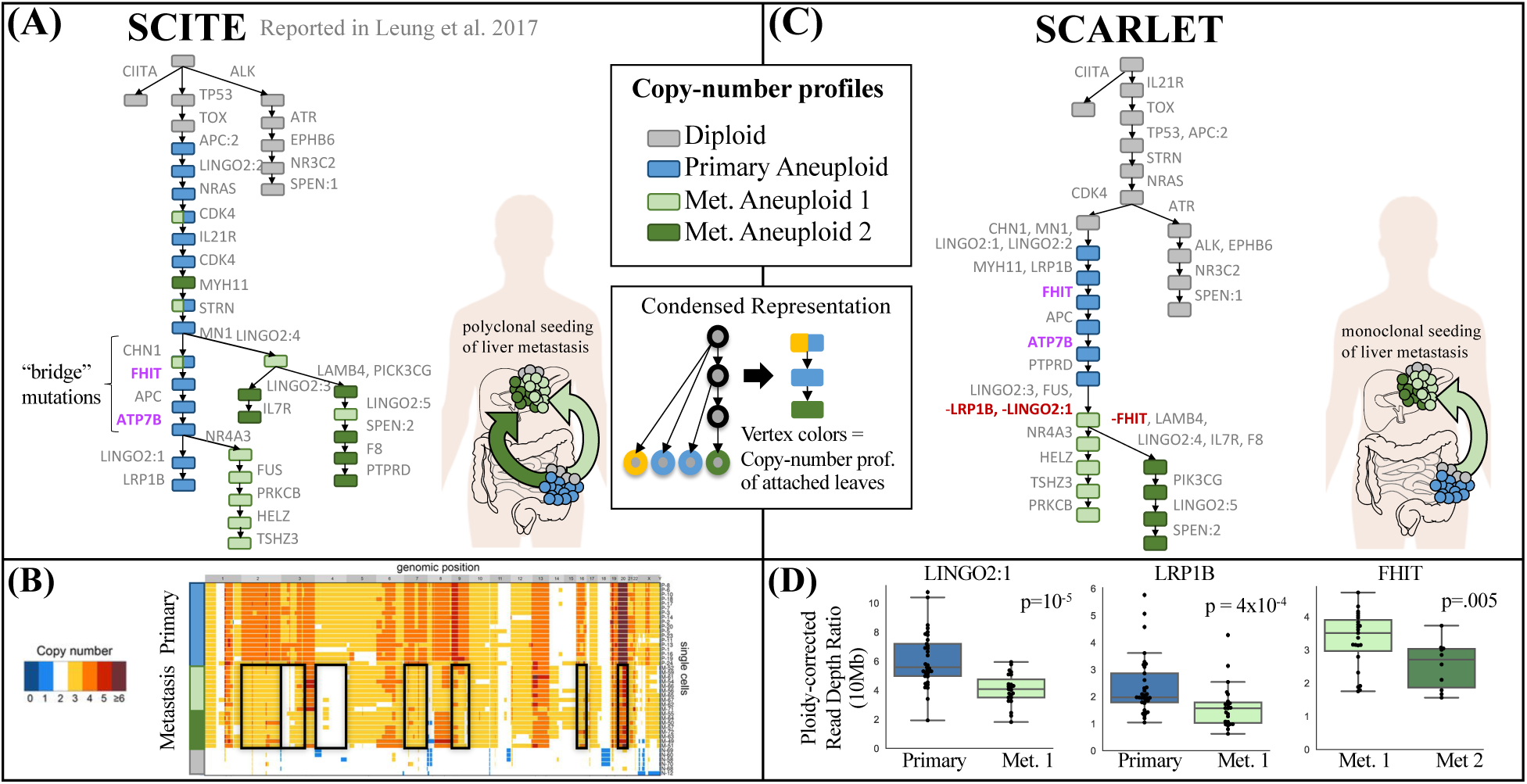
Scarlet infers a loss-supported phylogeny consistent with copy-number profiles from a metastatic colorectal cancer patient. (A) Published perfect phylogeny tree^36^ of 141 single cells from the primary colon tumor (blue) and 45 single-cells from the liver metastasis (green) of patient CRC2. Two distinct branches of metastatic cells — suggesting polyclonal seeding of the liver metastasis – are separated by four “bridge mutations” occurring in cells of the primary tumor. (B) Published copy-number profiles from DOP-PCR whole-genome sequencing of 42 single cells from both the primary tumor and metastasis of CRC2 (figure adapted from Leung et al.^36^). All metastatic cells share deletions of six chromosomes (black boxes), but are separated into two groups (light and dark green) by a small number of additional CNAs. (C) Scarlet infers a loss-supported phylogeny from the same data. The Scarlet tree has a single branch containing all metastatic cells – suggesting monoclonal seeding of the liver metastasis and consistent with the similar copy-number profiles of all metastatic cells. Scarlet identifies mutation losses (red) in `LINGO2`, `LRP1B`, and `FHIT`. (D) Significant decreases in read depths are observed at the loci of the three mutation losses identified by Scarlet.

We analyzed this dataset using Scarlet to see whether joint analysis of SNVs and CNAs data could help resolve the inconsistency between the tree derived from SNVs and the observed copy number profiles. We first derived four distinct copy-number profiles by hierarchical clustering of ploidy-corrected read-depth ratios from the targeted single-cell sequencing data. These copy-number profiles included an aneuploid profile for all primary cells (P), two different aneuploid profiles for metastatic cells (M1 and M2), and the profile of diploid cells (D); Leung et al.^36^ similarly derived four copy-number profiles from whole-genome sequencing of a different set of 42 cells from the same patient. Since four copy-number profiles is a small number to infer a tree using a copy-number evolution model, we instead ran Scarlet in the ‘Optimal CN Tree’ setting selecting the copy-number tree that produced the highest likelihood. Specifically, we ran Scarlet on all nine possible rooted copy-number trees with the root having the diploid profile (D), and internal vertices labeled by one of the three aneuploid copy profiles (P, M1, M2). For each copy-number tree, we derived the set ℒ of supported losses as the mutation loci that exhibited significant decreases in read depth (i.e., number of aligned sequencing reads). Additional details are in Section S2.4.

Scarlet constructed a tree (Fig. 4(C) and Fig. S1(B)) with a single clade containing all metastatic cells. This is consistent with the copy-number data, since the shared chromosomal losses could have have occurred once in a common ancestor of all metastatic cells. Moreover, this tree suggests that the liver metastasis was the product of *monoclonal seeding*; i.e., a single cell (or small group of cells) with the same somatic mutations migrated from the primary colon tumor to the metastasis and all metastatic cells descended from the founder cells present in this single migration. This result contradicts previous results^32, 36^ of a more complicated polyclonal seeding of the metastasis. The Scarlet tree contains three mutation losses: in genes `FHIT`, `LRP1B`, and `LINGO2`. Each of these losses is supported by a significant decrease in read depth (Fig. 4(D)), providing evidence that the loci containing these mutation were likely affected by deletions. Notably `FHIT` and `LRP1B` are located in fragile sites in the genome^42^, which are known regions of genomic instability. In addition, the loss of the mutation `LINGO2:1` in `LINGO2` is further supported by a shift in the variant allele frequency of another mutation, `LINGO2:2`, in the same gene. Specifically, the variant allele frequency of `LINGO2:2` is ≈1 in the metastatic cells (Fig. S1(A)), suggesting that this mutant allele is homozygous, consistent with a deletion or loss of heterozygosity event where the `LINGO2:1` mutations was lost.

We examined further the evidence for polyclonal seeding in the initial study of this patient. Leung et al.^36^ included a statistical analysis of the variant read counts of the four “bridge mutations”, `ATP7B`, `FHIT`, `APC` and `CHN1` that occurred between the first and second metastatic branches in the SCITE tree. This analysis showed that mutations in `ATP7B` and `FHIT` were present in a subset of primary tumor cells and in the second metastatic branch (detected in 10/13 and 13/13 cells respectively) while being absent in the second metastatic branch (detected in 1/15 and 1/15 cells respectively). Under the infinite sites model used by SCITE, mutation loss is not allowed and thus polyclonal seeding is necessary to explain the absence of these mutations. The same analysis found high uncertainty regarding the placement of mutations in `APC` and `CHN1` and thus these were not cited as evidence for polyclonal seeding.

The loss-supported model used by Scarlet provides an alternate explanation for the absence of `FHIT` and `ATP7B`. Scarlet identifies a supported mutation loss to explain the presence of the mutation in `FHIT` only in a subset of metastatic cells (M1). This loss is supported by a shift in read depth (*p* = 0.005) in the 10Mb region containing the locus (Fig 4D). Scarlet does not identify a supported mutation loss to similarly explain `ATP7B` as we did not observe a significant decrease in read depth for the corresponding locus (*p* = 0.34). However, this lack of a significant decrease in read depth at the `ATP7B` locus does not necessarily imply that there was no mutation loss. In particular, because targeted sequencing was performed for only 1000 genes, the copy number data is fairly low resolution and we calculated read depth in 10Mb bins. Thus, we may lack the statistical power to identify a shorter deletion, especially a deletion present in only the 10 metastatic cells with copy number profile M2. In summary, we argue that the sequencing data provides stronger evidence for the phylogeny constructed by Scarlet, which is consistent with both SNV and copy-number data, and supports a more parsimonious explanation of monoclonal seeding of the liver metastasis.

## 3 Discussion

Somatic mutations in tumors range across all genomic scales, from single-nucleotide variants (SNVs) through large copy-number aberrations (CNAs). To date, most methods for constructing phylogenies from single-cell DNA sequencing (scDNA-seq) data^21–24, 28–33^ used only SNVs, ignoring CNAs and thus throwing out important information for phylogenetic inference. Here, we introduced Scarlet, the first algorithm – to our knowledge – that uses measurements of *both* SNVs *and* CNAs to reconstruct tumor phylogenies from scDNA-seq data. Scarlet is based on a novel loss-supported evolutionary model, which constrains mutation losses to loci containing evidence of a CNA. By using the information about CNAs that is readily available in scDNA-seq data, the loss-supported model has less ambiguity in the phylogeny inference than the Dollo and finite sites models which allow mutation losses to occur anywhere on the tree. In scDNA-seq data, where there is often considerable uncertainty in the mutations present in each cell, this reduction in ambiguity enables more accurate phylogeny inference. On simulated scDNA-seq data, we find that Scarlet outperforms existing methods that do not utilize copy-number data. On targeted scDNA-seq data from a metastatic colorectal cancer patient, we showed that Scarlet found a phylogeny containing three mutation losses. Notably, Scarlet’s tree was both more consistent with the copy-number data and provided a simpler explanation of monoclonal seeding of the liver metastasis, compared to the more complex phylogenies reported previously^32, 36^. Thus, accurate modeling of mutations losses results in different conclusions regarding the migration patterns of metastasis.

There are a number of directions for future improvement. First, the current implementation of Scarlet either requires the copy-number tree in input or enumerates all possible copy-number trees and selects the maximum likelihood result. This approach is applicable when the number of distinct copy-number profiles is small; e.g., in the case of targeted scDNA-seq data^17, 43, 44^ where copy-number data typically is lower resolution. However, with higher-quality copy-number data, extensions to larger numbers of copy-number profiles is needed. One approach is to use copy-number evolution models^13, 37–39^ to identify a modest number of copy-number trees that summarize the uncertainty in the copy-number evolutionary history. Second, one could extend the loss-supported model into a unified evolutionary model for SNVs and CNAs. Indeed, the loss-supported model provides a natural framework to integrate SNVs directly with evolutionary models of CNAs. As single-cell sequencing technologies continue to improve, higher quality measurements of both SNVs and CNAs from the same sets of cells will become available. We anticipate that Scarlet and the loss-supported model will play a crucial role in the analysis of these data.

## 4 Methods

### 4.1 Loss-supported phylogeny model

We model the evolutionary history of a tumor as a rooted, directed phylogenetic tree *T* = (*V* (*T*), *E*(*T*)), whose vertex set *V* (*T*) = *L*(*T*) ∪ *I*(*T*) consists of a set *L*(*T*) of *n* leaves corresponding to *observed cells* and a set *I*(*T*) of inner vertices corresponding to *ancestral cells*. A directed edge (*v, w*) ∈ *E*(*T*) indicates that cell *v* is an ancestor of cell *w*. We do not directly observe *T* but rather we measure a set of phylogenetic markers for every observed cell *v* ∈ *L*(*T*). In the case where the markers are somatic single-nucleotide variants (SNV), the measurements correspond to a binary *mutation profile* **b**_*v*_ ∈ {0, 1}^*m*^ for each observed cell *v*, where *b*_*v,a*_ = 1 indicates that cell *v* has a somatic mutation at locus *a* and *b*_*v,a*_ = 0 indicates that cell *v* does not have a somatic mutation at locus *a*. We assume that the mutation profile **b**_*r*_ of the root *r* is 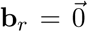 since the root represents the normal cell that preceded the tumor. We define the *mutation matrix* **B** = [**b**_*v*_]_*v*∈*L*(*T*)_ to be the matrix whose rows are the mutation profiles of leaves *v* ∈ *L*(*T*).

The problem of phylogenetic tree inference is to find a tree *T* and an *augmented mutation matrix* 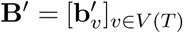 whose rows correspond to binary mutation profiles of the vertices of *T* and where the submatrix 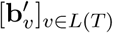 is equal to **B**. Since there are many possible trees that relate the observed cells, methods for phylogeny inference find *T* and **B**′ that best fit a specific evolutionary model.

The simplest evolutionary model for SNVs is the *infinite sites*, or *perfect phylogeny* model. In this model, each mutation is gained (0 → 1) at most once, and is never subsequently lost. A more general model the Dollo model allows mutations to be gained (0 → 1) at most once, but lost (1 → 0) multiple times.

Formally, the Dollo model is defined as follows.

#### Definition 1.

*A phylogenetic tree T is a* Dollo phylogeny *with respect to augmented mutation matrix* **B**^′^ *provided that for every locus a, there is at most one edge* (*v, w*) ∈ *E*(*T*) *such that* 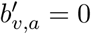 *and* 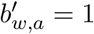.

In contrast to the perfect phylogeny model, under the Dollo model there are often multiple phylogenies that are consistent with input data (Fig. 1).

DNA sequencing data often contains contains additional information about the genomic locations where mutation losses are possible. Specifically, we assume that for each cell *v*, we also observe a copy-number profile **p**_*v*_ = [*p*_*v*,1_, …, *p*_*v,N*_] where *p*_*v,i*_ indicates the number of copies of genomic segment *i* in cell *v*. For simplicity, we label the unique copy-number profiles observed for all the cells by integers {1, …, *k*}, such that the vector **c** = [*c*_*v*_] represents the copy-number profile assignment *c*_*v*_ ∈ {1, …, *k*} of every cell *v*. The copy-number profiles of cells provide constraints on mutation losses. In particular, we allow mutation losses only at loci where an overlapping deletion or loss-of-heterozygosity (LOH) distinguishes the copy-number profiles. We record the information about the loci where losses are allowed in a collection ℒ of *supported loss sets*. For each pair *c, c*′ of distinct copy-number profiles we define the set ℒ (*c, c*′) ⊆ {1, …, *m*} of *supported losses* to be the set of all the mutation loci located in genomic regions with a decrease in copy number (indicating possible deletion or LOH) between *c* and *c*′. We define ℒ (*c, c*) = ø for all *c*. We denote the collection of supported losses as ℒ = {ℒ (*c, c*′): (*c, c*′) ∈ {1, …, *k*} × {1, …, *k*}}. We define a *loss-supported phylogeny* as a Dollo phylogeny where all mutation losses are supported.

#### Definition 2.

*Given copy number profiles* **c**′ = [*c*_*v*_]_*v*∈*V* (*T*)_ *and supported losses* ℒ, *a phylogenetic tree T is a* loss-supported phylogeny *with respect to augmented mutation matrix* **B**′ *provided that: (1) T is a Dollo phylogeny; (2) If* 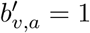 *and* 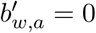 *for edge* (*v, w*) *and locus a then* 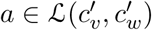.

The loss-supported phylogeny inference problem is to infer a loss-supported phylogeny *T* given a mutation matrix **B** and copy-number profile vector **c** that label the leaves of *T*, as well as a set ℒ of supported losses. However, this general problem has a major complication: the copy-number profiles of the ancestral cells are unknown. Without knowledge of ancestral copy-number profiles, the loss sets ℒ cannot be used to constrain mutation losses. Ideally, one might infer copy-number profiles of ancestral cells (e.g., using a copy-number evolution model^13, 37–39^) while *simultaneously* inferring a loss-supported phylogeny on the SNVs. The derivation of a score/likelihood for such joint model is not straightforward, and is left for future work. Instead, in the next section, we describe an algorithm that infers a loss-supported phylogeny by *refining* a *copy-number tree* given in input.

### 4.2 Loss-supported Refinement Problem

In this section, we introduce the *Loss-Supported Refinement (LSR) problem*, a special case of the loss-supported phylogeny inference problem, where we have additional information about the evolutionary relationships between copy-number profiles. In particular, we assume that we are given a *copy-number tree T* = (*V* (*T*), *E*(*T*)) and a copy-number profile vector **c** = [*c*_*v*_]_*v*∈*V* (*T*)_ for all vertices in *T*. As single-cell DNA sequencing data of SNVs typically measures copy-number profiles with low-resolution, this copy-number tree typically has many multifurcations. (i.e., unresolved ancestral vertices with more than two children). We use the mutation matrix **B** = [**b**_*v*_] for all *v* ∈ *L*(*T*) to refine vertices in *T*, which results in a *joint tree T*′ that reflects the evolutionary history of both the SNVs and CNAs. This sequential approach is inspired by an asymmetry between SNVs and CNAs in the loss-supported model: CNAs affect the observed state transitions of SNVs as deletions result in SNV loss, but SNVs do not result in changes in copy-number state.

The joint tree *T*′ is a *refinement*^45^ *of T*; i.e., *T* may be obtained by contracting edges in *T*′, according to the following definition.

#### Definition 3.

*A tree T*′ *is a refinement of a tree T provided L*(*T*′) = *L*(*T*) *and there exists a mapping γ*: *V* (*T*) → 2^*V* (*T*′)^ *satisfying the following conditions.*

1. *For all v* ∈ *V* (*T*), *γ*(*v*) *is a rooted subtree T*′ [*γ*(*v*)] *of T*′ *with root r*(*v*). (Contiguity)
2. *For all* (*v, w*) ∈ *E*(*T*), *there exists exactly one edge* (*p*(*r*(*w*)), *r*(*w*)) ∈ *E*(*T*′) *such that p*(*r*(*w*)) ∈ *γ*(*v*). (Edge consistency)
3. *For all v* ∈ *L*(*T*), *γ*(*v*) = {*v*}. (Leaf consistency)

We define the LSR problem as the problem of finding a refinement *T*′ of a copy-number tree *T* such that *T*^′^ is a loss-supported phylogeny.

#### Problem 1.

***Loss-Supported Refinement (LSR) problem***

*Given a copy-number tree T, a copy-number profile vector* **c** = [*c*_*v*_]_*v*∈*V* (*T*)_, *a mutation matrix* **B** = [**b**_*v*_]_*v*∈*L*(*T*)_, *and supported losses* ℒ, *find a refinement T*′ *of T, a copy-number profile vector* **c**′ = [*c*_*v*′_]_*v*′∈*V* (*T*′)_, *and an augmented mutation matrix* 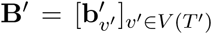 *with* 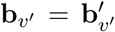 *for all v*′ ∈ *L*(*T*′), *such that*

1. 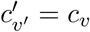 *for all v* ∈ *V* (*T*) *and v*′ ∈ *γ*(*v*), *and*
2. *T*′ *is a loss-supported phylogeny with respect to* **B**′, **c**′, *and* ℒ.

We provide four sufficient and necessary conditions for a solution *T*′, **c**′, **B**′ to the LSR problem.

#### Theorem 1.

*Given copy-number tree T, copy-number profile vector* **c**, *mutation matrix* **B**, *and supported losses* ℒ, *a refinement T*′ *of T, copy-number profile vector* **c**′, *and augmented mutation matrix* **B**′ *are a solution to the LSR problem if and only if*

1. *For all loci a, there exists exactly one edge* (*v*′, *w*′) ∈ *E*(*T*′) *with* 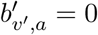 *and* 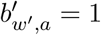; *And for all v* ∈ *V* (*T*):
2. *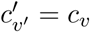 for all v*′ ∈ *γ*(*v*);
3. *If 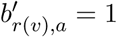 then 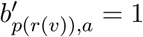 for all 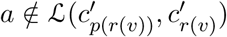;*
4. *There does not exist any edge* (*v*′, *w*′) ∈ *E*(*T*′ [*γ*(*v*)]) *with 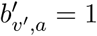 and 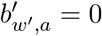.*

Note that Theorem 1 characterizes a solution *T*′ to the LSR problem as composed of *n* + *k* subtrees *T*′ [*γ*(*v*)] for *v* ∈ *V* (*T*) (Fig. 2). Moreover, conditions (1) and (4) imply that each of these subtrees *T*′ [*γ*(*v*)] is a perfect phylogeny with respect to submatrix **B**′ [*γ*(*v*)]. We use this structure to solve the LSR problem in the next section.

### 4.3 Solving the Loss-Supported Refinement problem

In this section, we derive an efficient algorithm to solve the LSR problem. This algorithm decomposes the LSR problem into *k* = |*I*(*T*)| instances – one for each copy-number profile – of the Incomplete Directed Perfect Phylogeny (IDP) problem^46^, using the characterization of LSR solutions given in Theorem 1. Specifically, Theorem 1 characterizes LSR solutions by giving a set of conditions on the set 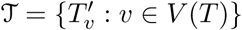 of subtrees of *T* defined by the refinement mapping *γ*, such that 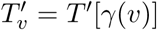. We design an algorithm to find a set 𝒯 of subtrees, mutation matrix **B**′, and copy-number profiles **c**′ that satisfy conditions (1)-(4) of Theorem 1. Using 𝒯 and **B**′, we then construct a refinement *T*′ such that 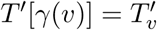 and 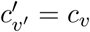 for all vertices 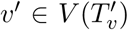. By conditions (1) and (4), each subtree 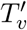 is a perfect phylogeny with respect to the corresponding mutation submatrix 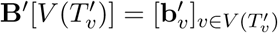. Moreover, by condition (3), 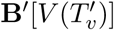 is constrained by the mutation profiles 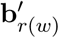 of the root *r*(*w*) of every descendent subtree 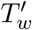 with (*v, w*) ∈ *E*(*T*). Thus, we recursively solve for 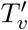 and 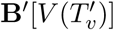 for every vertex *v* ∈ *V* (*T*) starting from the leaves *L*(*T*), as *L*(*T*) do not have any descendants and thus can be solved independently. The algorithm relies on three additional constraints on the solution *T*′, **c**′ and **B**′, described in the following lemma.

#### Lemma 1.

*If there exists a solution to the LSR problem for a given T*, **c, B**, ℒ, *then there exists a solution T*′, **c**′, **B**′ *that satisfies the following conditions.*

i. *For all* (*v, w*) ∈ *E*(*T*), *p*(*r*(*w*)) *is a leaf of subtree T*′ [*γ*(*v*)].
ii. *For all v* ∈ *V* (*T*)\{*r*}, *if 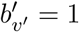 for all v*′ ∈ *L*(*T*′ [*γ*(*v*)]) *then 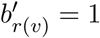.*
iii. *For all v* ∈ *V* (*T*) *and all loci a, 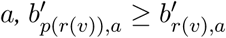.*

Our recursive algorithm is composed of a base and recursive step.

#### Base step

The base case determines 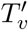 and 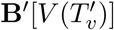 for leaf vertices *v* ∈ *L*(*T*). By Definition 3, *γ*(*v*) = {*v*} for any leaf in a refinement *T*′. Thus the subtree 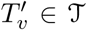 is composed of a single vertex *v*, with mutation profile 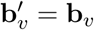 and copy-number profile 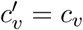.

#### Recursive step

The recursive step aims to find 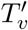 and 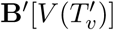 for internal vertices *v* ∈ *I*(*T*). We recursively solve for 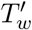 and 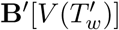 for every vertex *w* such that (*v, w*) ∈ *E*(*T*). By condition (i) of Lemma 1, 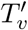 has a leaf for every (*v, w*) ∈ *E*(*T*), such that 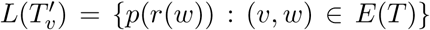. We know that 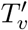 is a perfect phylogeny with respect to 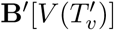 (condition (3)), and the structure of perfect phylogeny 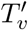 is determined when the mutation profiles 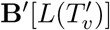 of the leaves are given^25^. Therefore, identifying 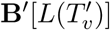 is sufficient to obtain 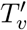 and 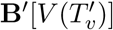.

We do not directly observe 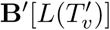, but there are two constraints on 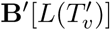 in addition to 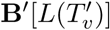 being a perfect phylogeny matrix. We summarize these two constraints on 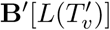 as a ternary matrix 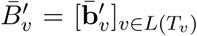 such that 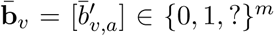. The first constraint is provided by condition (2) of Theorem 1 and condition (iii) of Lemma 1 which fix the values for some entries of 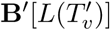 such that 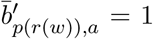 when 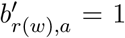 and 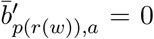 when 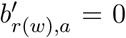 and *a* ∈ ℒ (*c*_*v*_, *c*_*w*_). The second constraint is provided by condition (4) and we meet this constraint by further setting some of the previously non-fixed entries in 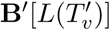 to minimize the total number of mutation gains in 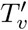. A mutation gain is present in 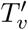 only if there are vertices *v*′, *w*′ ∈ *L*(*T*_*v*_) such that 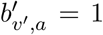 and 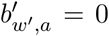. To achieve the minimum number of mutation gains, we thus maximize the number of all-zero and all-one columns of 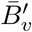: we set to 0 any previously undetermined entries 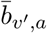 for columns of 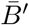 that only have ‘0’ (‘1’, resp.) entries (setting of 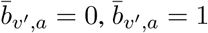 resp.). At last, we set any remaining undetermined entry of 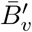 to be ‘?’.

Finally, we aim to find **B**′ [*L*(*T*_*v*_)] by filling the ‘?’ entries of 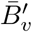. More specifically, given 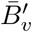, we seek **B**′ [*L*(*T*_*v*_)] such that if 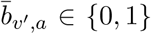 then 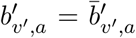 for all mutations *a* and **B**′ [*L*(*T*_*v*_)] is a perfect phylogeny matrix. This problem is known as the Incomplete Directed Perfect Phylogeny (IDP) problem and has been shown to be solvable in *O*(*n*^2^*m*) time^46^. In our case *n* = |*L*(*T*_*v*_)| = *d*_*v*_ where *d*_*v*_ is the out-degree of vertex *v* in *T*. Solving an instance of the IDP problem yields a perfect phylogeny mutation matrix **B**′ [*L*(*T*_*v*_)], which in turn determines the perfect phylogeny tree 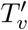 and mutation matrix 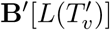.

### 4.4 Maximum Likelihood Loss-supported Refinement Problem

The LSR problem assumes that the mutation matrix **B** is error-free. In practice, we do not observe this mutation matrix **B**, but instead we observe read counts from a sequencing experiment. Specifically, we measure a variant read count matrix **X** = [**x**_*v*_]_*v*∈*L*(*T*)_ and a total read count matrix **Y** = [**y**_*v*_]_*v*∈*L*(*T*)_, where *x*_*v,a*_ ∈ ℕ is the number of variant reads at locus *a* in cell *v* and *y*_*v,a*_ ∈ ℕ is the total number of reads. Whole-genome amplification^18^, which typically precedes single-cell DNA sequencing, introduces a considerable amount of error into these read count matrices. Specifically, single-cell sequencing SNV data has high rates of false negative errors (i.e., *x*_*v,a*_ = 0 when *b*_*v,a*_ = 1) and missing data (i.e., *y*_*v,a*_ = 0). In addition, sequencing and whole-genome amplification introduce false positive errors (i.e., *x*_*v,a*_ > 0 when *b*_*v,a*_ = 0) as well. Most existing methods^21, 22, 24, 28, 31–33^ for single-cell phylogeny inference discretize read counts into an observed mutation matrix 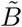, using either two or three genotypes in addition to missing data (i.e, 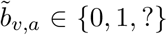 or 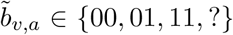). However, discretizing the mutation data loses information about the likelihood of errors. For example, a locus with a single variant read is far more likely to be a false positive error than a locus with hundreds of variant reads, but a discretized mutation matrix does not distinguish between these cases.

Here, we adopt a maximum-likelihood approach that models the observed variant and total read counts. A similar approach was used in SciΦ^23^ with the infinite sites model for SNVs. Our approach aims to find the mutation matrix **B**\* = argmax Pr(**X** | **Y, B**); i.e., the mutation matrix that admits a solution *T*′, **B**′, **c**′ to the LSR problem and maximizes the likelihood of the observed variant read counts **X** given the total read counts **Y**. This formulation is not specific to a particular likelihood model for read counts but does assume that variant read counts **X** are independent of each other across cells and loci given **Y** and **B** – i.e, a likelihood of the form 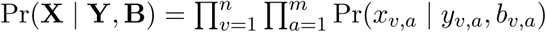 Let ℬ_*T*,**c**, ℒ_ be the set of mutation matrices **B** such that there exists a solution *T*′, **c**′, **B**′ to the LSR problem given *T*, **c**, and ℒ. We formulate the problem as follows.

#### Problem 2.

***Maximum Likelihood Loss-Supported Refinement (ML-LSR) problem***

*Given variant read counts* **X** = [**x**_*v*_]_*v*∈*L*(*T*)_, *total read counts* **Y** = [**y**_*v*_]_*v*∈*L*(*T*)_, *copy-number tree T, copy-number profile vector* **c** = [*c*_*v*_]_*v*∈*V* (*T*)_, *and supported losses* ℒ, *find the mutation matrix* **B**\* ∈ ℬ_*T*,**c**, ℒ_ *that maximizes* Pr(**X** | **Y, B**).

We show the ML-LSR is NP-hard by reduction from the Minimum Flip Problem^47^ (Section S3.3). Since current datasets have mutation matrices with hundreds–thousands of cells, we derive an algorithm in the next section that finds an approximate solution to the ML-LSR problem by subdividing the ML-LSR problem into *k* instances of the maximum likelihood Incomplete Directed Perfect Phylogeny problem.

### 4.5 Scarlet Algorithm for Maximum-Likelihood Loss-Supported Refinement Problem

We introduce Scarlet (<u>S</u>ingle-<u>C</u>ell <u>A</u>lgorithm for <u>R</u>econstructing the <u>L</u>oss-supported <u>E</u>volution of <u>T</u>umors), an algorithm to find a loss-supported phylogeny *T*′ from single-cell DNA sequencing data. Scarlet aims to solve the ML-LSR problem by finding the maximum likelihood mutation matrix **B**^*^ such that there exists a solution *T*′, **c**′, **B**′ to the LSR problem where **B**′ is an augmentation of **B**\* (i.e., **B**\* = **B**′ [*L*(*T*)] = **B**′ [*L*(*T*′)]). We proceed here by finding **B**′, which gives us **B**\*. In Section 2.3, we showed that finding **B**′ for the LSR problem decomposes into a set of IDP instances if we know the mutation profiles 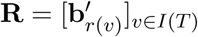 of the roots of subtrees 𝒯. Given **B**, we inferred **R** recursively, starting with the leaves *L*(*T*) whose mutation profiles are known. In the ML-LSR, however, we are not given **B**. Therefore, Scarlet uses a heuristic to find **B**\* where we first compute maximum likelihood mutation profiles **R**\* of the roots, and then solve a set of instances of a maximum likelihood IDP (ML-IDP) problem given **R**\*. We compute *R*\* by marginalizing over possible mutation profiles for each mutation (Section S2.2). As ML-IDP input, we define a ternary matrix 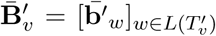 for each vertex *v* ∈ *V* (*T*). For *v* ∈ *I*(*T*), we define 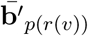 as in Section 4.3 given the mutation profile 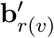 of the root *r*(*v*) of 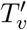. For *v* ∈ *L*(*T*), we have that 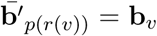 but unlike in the LSR problem, we are not given the mutation profile **b**_*v*_ in the ML-LSR problem. Instead, we are able to compute the likelihood of **b**_*v*_ as in Equation 1. As such, finding **B**\* is equivalent to find the maximum likelihood submatrices {**B**\*[{*v*: *c*_*w*_ = *c*_*v*_}]: *v* ∈ *I*(*T*)} such that 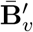 admits an incomplete directed perfect phylogeny. We show in Section S2.3 how to compute these maximum likelihood submatrices using an integer-linear programming (ILP) formulation.

This heuristic is not guaranteed to find the overall maximum likelihood solution **B**\* as there may be cases where **B**\* does not admit a solution with the maximum likelihood set of roots **R**\*. However, we showed in the Section 2.2 that Scarlet is both accurate and fast in practice. Scarlet can be used with any likelihood model that assumes conditional independence between variant read counts given the mutation matrix and total read counts as described in the previous section. In this work, we used a beta-binomial model for variant read counts, similar to the one used by SciΦ^23^, which accounts for overdispersion due to whole-genome amplification as well as sequencing error. Additional details are in Section S2.1.

## Acknowledgements

This work is supported by a US National Institutes of Health (NIH) grants R01HG007069 and U24CA211000, US National Science Foundation (NSF) CAREER Award (CCF-1053753) and Chan Zuckerberg Initiative DAF grants 2018-182608 to BJR.

## Code and Data Availability

Scarlet software, simulated data, and processed CRC2 data are available at github.com/raphael-group/scarlet. Original CRC2 data was downloaded from NCBI Sequence Read Archive (SRA; https://www.ncbi.nlm.nih.gov/sra) under accession number SRP074289.

## References

[1] Burrell, R. A., McGranahan, N., Bartek, J. & Swanton, C. The causes and consequences of genetic heterogeneity in cancer evolution. Nature 501, 338–345 (2013).

[2] Tabassum, D. P. & Polyak, K. Tumorigenesis: it takes a village. Nature Reviews Cancer 15, 473–483 (2015).

[3] Jiao, W., Vembu, S., Deshwar, A. G., Stein, L. & Morris, Q. Inferring clonal evolution of tumors from single nucleotide somatic mutations. BMC bioinformatics 15, 35 (2014).

[4] El-Kebir, M., Oesper, L., Acheson-Field, H. & Raphael, B. J. Reconstruction of clonal trees and tumor composition from multi-sample sequencing data. Bioinformatics 31, i62–i70 (2015).

[5] Malikic, S., McPherson, A. W., Donmez, N. & Sahinalp, C. S. Clonality inference in multiple tumor samples using phylogeny. Bioinformatics 31, 1349–1356 (2015).

[6] Popic, V. et al. Fast and scalable inference of multi-sample cancer lineages. Genome biology 16, 91 (2015).

[7] Deshwar, A. G. et al. Phylowgs: reconstructing subclonal composition and evolution from whole-genome sequencing of tumors. Genome biology 16, 35 (2015).

[8] El-Kebir, M., Satas, G., Oesper, L. & Raphael, B. J. Inferring the mutational history of a tumor using multi-state perfect phylogeny mixtures. Cell systems 3, 43–53 (2016).

[9] Jiang, Y., Qiu, Y., Minn, A. J. & Zhang, N. R. Assessing intratumor heterogeneity and tracking longitudinal and spatial clonal evolutionary history by next-generation sequencing. Proceedings of the National Academy of Sciences 113, E5528–E5537 (2016).

[10] Alves, J. M., Prieto, T. & Posada, D. Multiregional tumor trees are not phylogenies. Trends in cancer 3, 546–550 (2017).

[11] Satas, G. & Raphael, B. J. Tumor phylogeny inference using tree-constrained importance sampling. Bioinformatics 33, i152–i160 (2017).

[12] Pradhan, D. & El-Kebir, M. On the non-uniqueness of solutions to the perfect phylogeny mixture problem. In RECOMB International conference on Comparative Genomics, 277–293 (Springer, 2018).

[13] Zaccaria, S., El-Kebir, M., Klau, G. W. & Raphael, B. J. Phylogenetic copy-number factorization of multiple tumor samples. Journal of Computational Biology 25, 689–708 (2018).

[14] Miura, S. et al. Power and pitfalls of computational methods for inferring clone phylogenies and mutation orders from bulk sequencing data. bioRxiv 697318 (2019).

[15] Myers, M. A., Satas, G. & Raphael, B. J. Calder: Inferring phylogenetic trees from longitudinal tumor samples. Cell systems (2019).

[16] 10X Genomics. Assessing tumor heterogeneity with single cell cnv. https://www.10xgenomics.com/solutions/single-cell-cnv. Accessed: 2019-Nov-05.

[17] Mission Bio. Copy number variants and single nucleotide variants simultaneously detected in single cells. https://missionbio.com/cnv_application_note. Accessed: 2019-Nov-05.

[18] Gawad, C., Koh, W. & Quake, S. R. Single-cell genome sequencing: current state of the science. Nature Reviews Genetics 17, 175 (2016).

[19] Zahn, H. et al. Scalable whole-genome single-cell library preparation without preamplification. Nature methods 14, 167 (2017).

[20] Navin, N. E. The first five years of single-cell cancer genomics and beyond. Genome research 25, 1499–1507 (2015).

[21] Jahn, K., Kuipers, J. & Beerenwinkel, N. Tree inference for single-cell data. Genome biology 17, 86 (2016).

[22] Ross, E. M. & Markowetz, F. Onconem: inferring tumor evolution from single-cell sequencing data. Genome biology 17, 69 (2016).

[23] Singer, J., Kuipers, J., Jahn, K. & Beerenwinkel, N. Single-cell mutation identification via phylogenetic inference. Nature communications 9, 5144 (2018).

[24] Malikic, S., Jahn, K., Kuipers, J., Sahinalp, S. C. & Beerenwinkel, N. Integrative inference of subclonal tumour evolution from single-cell and bulk sequencing data. Nature communications 10, 2750 (2019).

[25] Gusfield, D. Efficient algorithms for inferring evolutionary trees. Networks 21, 19–28 (1991).

[26] Taylor, A. M. et al. Genomic and functional approaches to understanding cancer aneuploidy. Cancer cell 33, 676–689 (2018).

[27] Bielski, C. M. et al. Genome doubling shapes the evolution and prognosis of advanced cancers. Nature genetics 50, 1189 (2018).

[28] El-Kebir, M. Sphyr: tumor phylogeny estimation from single-cell sequencing data under loss and error. Bioinformatics 34, i671–i679 (2018).

[29] Ciccolella, S. et al. Inferring cancer progression from single cell sequencing while allowing loss of mutations. bioRxiv 268243 (2018).

[30] McPherson, A. et al. Divergent modes of clonal spread and intraperitoneal mixing in high-grade serous ovarian cancer. Nature genetics 48, 758 (2016).

[31] Zafar, H., Tzen, A., Navin, N., Chen, K. & Nakhleh, L. Sifit: inferring tumor trees from single-cell sequencing data under finite-sites models. Genome biology 18, 178 (2017).

[32] Zafar, H., Navin, N., Chen, K. & Nakhleh, L. Siclonefit: Bayesian inference of population structure, genotype, and phylogeny of tumor clones from single-cell genome sequencing data. Genome Research (2019).

[33] Malikic, S. et al. Phiscs: a combinatorial approach for subperfect tumor phylogeny reconstruction via integrative use of single-cell and bulk sequencing data. Genome Research (2019).

[34] Dollo, L. The laws of evolution. Bull. Soc. Bel. Geol. Paleontol 7, 164–166 (1893).

[35] Kuipers, J., Jahn, K., Raphael, B. J. & Beerenwinkel, N. Single-cell sequencing data reveal widespread recurrence and loss of mutational hits in the life histories of tumors. Genome research 27, 1885–1894 (2017).

[36] Leung, M. L. et al. Single cell dna sequencing reveals a late-dissemination model in metastatic colorectal cancer. Genome research gr–209973 (2017).

[37] Schwarz, R. F. et al. Phylogenetic quantification of intra-tumour heterogeneity. PLoS computational biology 10, e1003535 (2014).

[38] Chowdhury, S. A. et al. Inferring models of multiscale copy number evolution for single-tumor phylogenetics. Bioinformatics 31, i258–i267 (2015).

[39] El-Kebir, M. et al. Complexity and algorithms for copy-number evolution problems. Algorithms for Molecular Biology 12, 13 (2017).

[40] Wang, Y. et al. Clonal evolution in breast cancer revealed by single nucleus genome sequencing. Nature 512, 155 (2014).

[41] Govek, K., Sikes, C. & Oesper, L. A consensus approach to infer tumor evolutionary histories. In Proceedings of the 2018 ACM International Conference on Bioinformatics, Computational Biology, and Health Informatics, 63–72 (ACM, 2018).

[42] Smith, D. I., Zhu, Y., McAvoy, S. & Kuhn, R. Common fragile sites, extremely large genes, neural development and cancer. Cancer letters 232, 48–57 (2006).

[43] Xu, X. et al. Single-cell exome sequencing reveals single-nucleotide mutation characteristics of a kidney tumor. Cell 148, 886–895 (2012).

[44] Leung, M. L. et al. Highly multiplexed targeted dna sequencing from single nuclei. Nature protocols 11, 214 (2016).

[45] Wu, T., Moulton, V. & Steel, M. Refining phylogenetic trees given additional data: An algorithm based on parsimony. IEEE/ACM Transactions on Computational Biology and Bioinformatics (TCBB) 6, 118–125 (2009).

[46] Pe’er, I., Pupko, T., Shamir, R. & Sharan, R. Incomplete directed perfect phylogeny. SIAM Journal on Computing 33, 590–607 (2004).

[47] Chen, D., Eulenstein, O., Fernandez-Baca, D. & Sanderson, M. Minimum-flip supertrees: complexity and algorithms. IEEE/ACM Transactions on Computational Biology and Bioinformatics (TCBB) 3, 165–173 (2006).

