## Supplementary Materials for "Single-cell tumor phylogeny inference with copy-number constrained mutation losses"

**4 References** **9**

### 1 Supplementary Tables and Figures

| Algorithm | Publication | Evolutionary Model | Input Format | Additional Data |
|---|---|---|---|---|
| OncoNem | Ross and Markowetz [2016] | Infinite Sites | Binary | |
| SCITE | Jahn et al. [2016] | Infinite Sites | Binary or Ternary | |
| B-SCITE | Malikic et al. [2019a] | Infinite Sites | Binary | Bulk Sequencing |
| SciPhi | Singer et al. [2018] | Infinite Sites | Read Counts | |
| SASC | Ciccolella et al. [2018] | Dollo | Binary | |
| SPhyR | El-Kebir [2018] | Dollo | Binary | |
| SiFit | Zafar et al. [2017] | Finite Sites | Binary or Ternary | |
| SiCloneFit | Zafar et al. [2019] | Finite Sites | Binary or Ternary | |
| PhiSCS | Malikic et al. [2019b] | Finite Sites | Binary | Bulk Sequencing |
| SCARLET | | Loss-Supported | Read Counts | Copy-number |

**Table 1:** Single-cell phylogenetic reconstruction algorithms

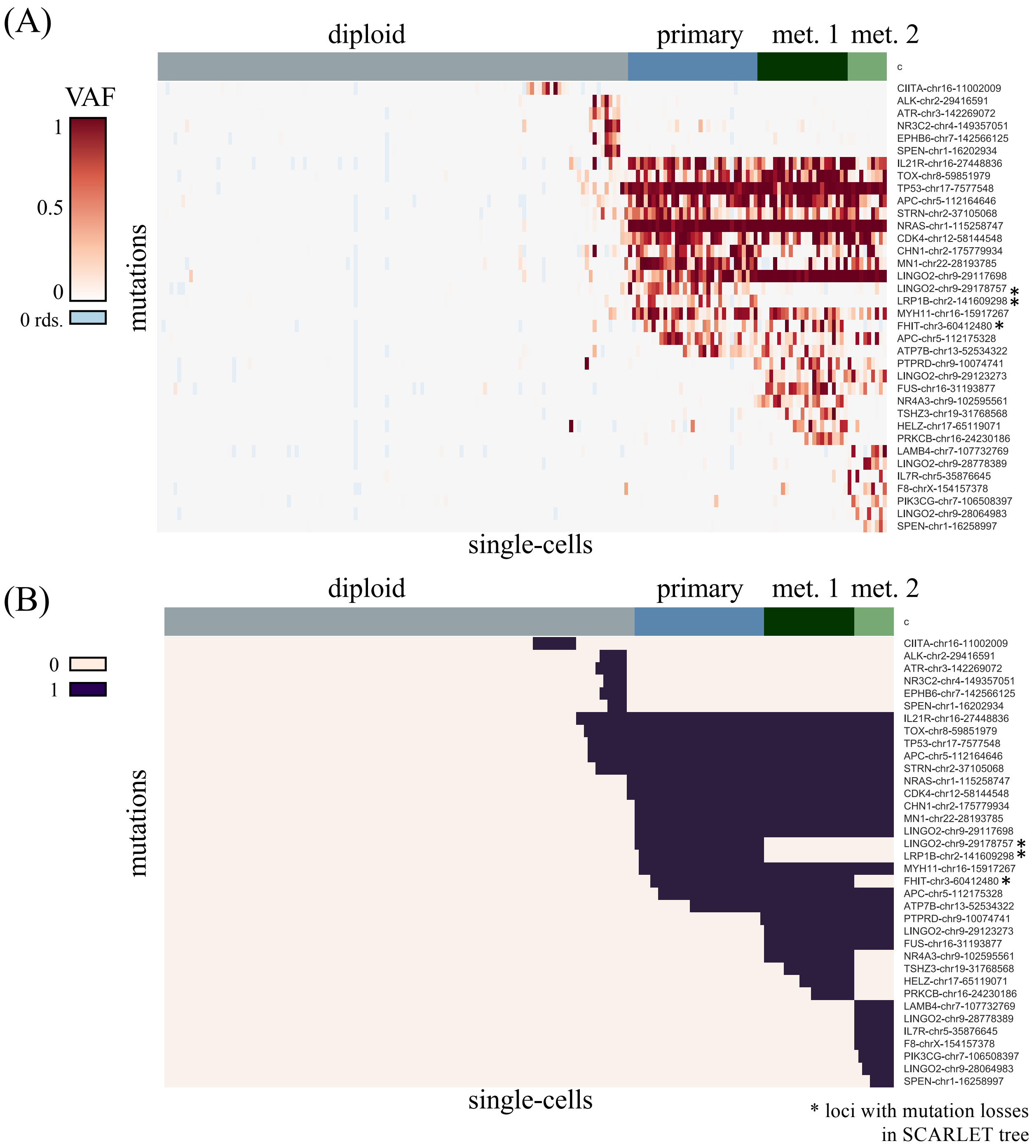

**Figure 1: CRC2 Data and SCARLET Mutation Matrix.** (A) Variant allele frequencies for 36 single-nucleotide variants detected in CRC2. (B) Maximum likeihood mutation matrix identified by SCARLET. Loci in which SCARLET detected a mutation loss are marked with an *.

### 2 Supplementary Methods

#### 2.1 Solving ML-LSR Problem: Likelihood model

We use a probabilistic model to evaluate mutation profiles using observed read counts. If mutation $a$ is absent in cell $v$ (i.e., $b_{v,a} = 0$), then the probability of observing a variant read corresponds to the per-nucleotide rate of sequencing error $\epsilon$. For Illumina sequencing reads, we use $\epsilon = 0.001$ **[CITE]**. If mutation $a$ is present in cell $v$ (i.e., $b_{v,a} = 1$), then we model the variant counts at a locus using a beta-binomial distribution (similarly to the model previously used by SciΦ Singer et al. [2018]). We estimate parameters $\alpha$ and $\beta$ empirically from the distribution of heterozygous germline single-nucleotide polymorphisms (SNPs) in the data. We thus define the data likelihood for observing $x_{v,a}$ variant reads at locus $a$ in cell $v$ as follows,

$$\Pr(x_{v,a} \mid y_{v,a}, b_{v,a}) = \begin{cases} \text{Beta-Binomial}(x_{v,a} \mid n = y_{v,a}, \alpha, \beta) & \text{if } b_{v,a} = 1, \\ \text{Binomial}(x_{v,a} \mid n = y_{v,a}, p = \epsilon) & \text{if } b_{v,a} = 0. \end{cases} \tag{1}$$

We assume variant read counts are independent across variants given $Y$ and $\mathbf{B}$. This gives us the following total likelihood:

$$\Pr(X \mid Y, B) = \prod_{v=1}^{n} \prod_{a=1}^{m} \Pr(x_{v,a} \mid y_{v,a}, b_{v,a}). \tag{2}$$

#### 2.2 Solving ML-LSR Problem: Finding max-likelihood subtree roots

As the first step of SCARLET, we find the maximum likelihood subtree roots $R$ for all internal subtrees. We must select a set of subtree roots $R$ such that there exists a loss-supported refinement $T'$, $\mathbf{c}'$, $\mathbf{B}'$ with subtree roots $R$. This constrains the possible mutation profiles of the roots. Specifically, by the Definition of a loss-supported phylogeny or each locus $a$, mutation $a$ is gained at most once in $T'$. Let subtree $T_a$ of $T$ correspond to the set of vertices $v$ for which which mutation $a$ is present in subtree $T[\gamma(v)]$. $R$ is a valid mutation state assignment for roots $R$ provided for each locus $a$, (1) that there exists a subtree $S_a$ of $T$ such that for all $v \in V(T)$, $r_{v,a} = 1$ if $v \in V(S_a)$ and $v$ is not the root of $S_a$ and $r_{v,a} = 0$ otherwise; and (2) for any edge $(v, w) \in T$ such that $v \in S_a$ and $w \notin S_a$, $a \in \mathcal{L}(c_v, c_w)$. Any valid $R$ uniquely defines a subtree $S_a$ for each locus $a$. Roots $R$ admit a mutation profile $\mathbf{b}_a$ provided that $\mathbf{b}_a$ satisfies the following.

1. If $v \notin S_a$ then mutation $a$ is absent in all cells $v'$ such that $c_{v'} = c_v$.
2. If $v \in S_a$ and $v$ is not the root of $S_a$ then mutation $a$ is present in all cells $v'$ such that $c_{v'} = c_v$.
3. If $v \in S_a$ and $v$ is the root of $S_a$ then mutation $a$ is either present or absent, as mutation $a$ occurred in $T'_v$

Let $\beta_R = \{\mathbf{b}_a : R \text{ admits } \mathbf{b}_a\}$ be the set of mutation profile for mutation $a$ admitted by roots $R$. The likelihood given a root assignment $R$ is computed by marginalizing over admitted mutation profiles, as follows.

$$\Pr(\mathbf{X} \mid \mathbf{Y}, \mathbf{R}) = \prod_{a=1}^{m} \prod_{\mathbf{b}_a \in \beta_R} \Pr(\mathbf{x}_a \mid \mathbf{y}_a, \mathbf{b}_a) = \prod_{a=1}^{m} \prod_{v=1}^{n} Pr(x_{v',a} \mid y_{v',a}, R) \tag{3}$$

such that

$$Pr(x_{v',a} \mid y_{v',a}, R) = \begin{cases} \Pr(x_{v',a} \mid y_{v',a}, b_{v',a} = 0) & \text{if } c_{v'} = c_v \text{ and } v \notin S_a \\ \Pr(x_{v',a} \mid y_{v',a}, b_{v',a} = 1) & \text{if } c_{v'} = c_v, v \in S_a \text{ and } v \text{ is not root of } S_a \\ \frac{1}{2}\Pr(x_{v',a} \mid y_{v',a}, b_{v',a} = 1) & \\ \quad +\frac{1}{2}\Pr(x_{v',a} \mid y_{v',a}, b_{v',a} = 0) & \text{otherwise} \end{cases} \tag{4}$$

We thus find $R^*$ by enumerating valid mutation state assignments for roots for each mutation locus $a$, then computing the maximum likelihood as above.

##### 2.3 Solving ML-LSR Problem: Integer Linear Program (ILP) to solve maximum likelihood IDP

We define a cost-function

$$C_{p,a} = \begin{cases} \log\Pr(x_{v,a} \mid y_{v,a}, b_{v,a} = 1) - \log\Pr(x_{v,a} \mid y_{v,a}, b_{v,a} = 0) & \text{if } v \text{ is observed,} \\ M & \text{if } v \text{ is not observed and } \bar{b}_{v,a} = 0 \\ -M & \text{if } v \text{ is not observed and } \bar{b}_{v,a} = 0 \\ 0 & \text{if } v \text{ is not observed and } \bar{b}_{v,a} = ?. \end{cases} \tag{5}$$

This gives us the following ILP. We introduce a set of auxiliary variables $F, G, H$ to enforce the three gametes condition, where $F_{a,b}$, $G_{a,b}$ and $H_{a,b}$ indicate that a pair of columns $a, b$ show the $(1, 1)$, $(0, 1)$ and $(1, 0)$ gametes respectively.

$$\begin{aligned}
&\text{maximize} \sum_{p,a} C_{p,a} \cdot B_{p,a} \\
&\text{subject to} \\
&\quad F_{a,b} + G_{a,b} + H_{a,b} \leq 2 && \text{for all } a, b \\
&\quad F_{p,a,b} \geq B_{p,a} + B_{p,b} - 1 && \text{for all } a, b, p \\
&\quad F_{p,a,b} \leq B_{p,a} && \text{for all } a, b, p \\
&\quad F_{p,a,b} \leq B_{p,a} && \text{for all } a, b, p \\
&\quad F_{a,b} \leq \sum_i F_{p,a,b} && \text{for all } a, b \\
&\quad F_{a,b} \geq F_{p,a,b} && \text{for all } a, b, p \\
&\quad G_{p,a,b} \geq (1 - B_{p,a}) + B_{b,i} - 1 && \text{for all } a, b, p \\
&\quad G_{p,a,b} \leq (1 - B_{p,a}) && \text{for all } a, b, p \\
&\quad G_{p,a,b} \leq B_{p,b} && \text{for all } a, b, p \\
&\quad G_{a,b} \leq \sum_i G_{p,a,b} && \text{for all } a, b \\
&\quad G_{a,b} \geq G_{p,a,b} && \text{for all } a, b, p \\
&\quad H_{p,a,b} \geq B_{p,a} + (1 - B_{p,b}) - 1 && \text{for all } a, b, p \\
&\quad H_{p,a,b} \leq B_{p,a} && \text{for all } a, b, p \\
&\quad H_{p,a,b} \leq (1 - B_{p,b}) && \text{for all } a, b, p \\
&\quad H_{a,b} \leq \sum_i H_{p,a,b} && \text{for all } a, b \\
&\quad H_{a,b} \geq H_{p,a,b} && \text{for all } a, b, p \\
&\quad B_{p,a} \in \{0, 1\} && \text{for all } a, p
\end{aligned}$$

| | |
|---|---|
| $F_{p,a,b} \in \{0,1\}$ | for all $a, b, p$ |
| $G_{p,a,b} \in \{0,1\}$ | for all $a, b, p$ |
| $H_{p,a,b} \in \{0,1\}$ | for all $a, b, p$ |
| $F_{a,b} \in \{0,1\}$ | for all $a, b$ |
| $G_{a,b} \in \{0,1\}$ | for all $a, b$ |
| $H_{a,b} \in \{0,1\}$ | for all $a, b$ |

##### 2.4 Simulation Details

We simulate 50 trees with 20 mutations and 4 copy-states. We do not directly model copy-number evolution in our simulation. Instead, for each edge between cells $p, q$ with different copy-number states $i, j$, we simulate the set of supported losses $\mathcal{L}_{i,j}$ by selecting a subset of loci of size $|\mathcal{L}_{i,j}| \sim$ Poisson$(0.2 * m)$. A mutation loss $-a$ is introduced on an edge with probability $0.5$ provided that the source cell $p$ contains mutation $a$, $a \in \mathcal{L}_{i,j}$. The resulting trees have between 1-8 mutation losses. In no instances was the same mutation lost multiple times on the same tree. Thus, all 50 trees respect the $k - Dollo$ model with $k = 1$. From each tree, we sample 100 cells and simulate sequencing error as follows. For each locus $a$ in cell $p$, we generate a variant read count $v_{p,a}$ and a total read count $r_{p,a}$. We simulate data with an allelic dropout rate of $d = 0.15$. With probability $d^2$, both alleles drop out, i.e., $v_{p,a} = r_{p,a} = 0$. Otherwise, the total number of reads is $r_{p,a} \sim$ Poisson$(100)$. If the variant is absent, $v_{p,a} \sim$ Binomial$(t_{p,a}, \epsilon)$ where $\epsilon = 0.001$ models the sequencing error rate. If the variant is present we model the overdispersion in the variant read distribution resulting from whole-genome amplification using a Beta-Binomial model. Let $f_{p,a} \sim \max\{\text{Beta}(\alpha, \alpha), \epsilon\}$ (with $\alpha = 0.25$ in order to obtain a dropout rate of $d \approx 0.15$) and $v_{p,a} \sim$ Binomial$(t_{p,a}, f_{p,a})$.

SCITE and SPhyR and SiFit were run with default parameters. In the case where SCITE returned multiple solutions, we selected the first solution that was returned. SPhyR was run with a maximum number $k = 1$ of mutation losses per locus. All algorithms were given the true allelic dropout and sequencing error rates. As SciPhi is designed for variant calling from a large set of putative variants, and thus has filters to remove false positives, we found that the default parameters removed a lot of the variants present in the tree. The parameters for these filters were (1) `cwm` the number of cells that show the mutation, (2) `nmc` the number of cells that must have a minimum alternate read count of (3) `ms`. As all variants including in the simulation are present in at least one cell, we set all three of these parameters to be 0, so as to not pre-filter out any variants.

##### 2.5 Copy-number analysis of CRC2

To assign cells to copy-number states for CRC2, we first binned reads at 10Mb. We next performed hierarchical clustering on cells. A cut-off for hierarchical clustering was selected to yield 4 clusters, the observed number of copy-number clones that was identified in the copy-number analysis performed by Leung et al. [2017]. Supported losses were inferred by looking for significant shifts in read-depth in the bin containing the variant of interest using a Wilcoxon rank-sum test and a p-value threshold of $p = 0.1$.

#### 3 Proofs

##### 3.1 Proof of Theorem 1

**Theorem 1.** *Given copy-number tree $T$, copy-number state vector* $\mathbf{c}$*, mutation matrix $B$ and supported losses $\mathcal{L}$, a phylogeny $T'$, copy-number state vector $c'$ and augmented mutation matrix $B'$ are a solution to* *the LSR problem if and only if $T'$ is a refinement of $T$ and for all $v \in V(T)$*

1. *, $c'_{v'} = c_v$ for all $v' \in \gamma(v)$;*
2. *$T'[\gamma(v)]$ is a perfect phylogeny with respect to $B'[\gamma(v)]$;*
3. *If $b'_{r(v),a} = 1$ then $b'_{p(r(v)),a} = 1$ for all $a \notin \mathcal{L}(c'_{p(r(v))}, c'_{r(v)})$;*
4. *For all mutations $a$, there exists exactly one edge $(v', w') \in E(T')$ with $b'_{v',a} = 0$ and $b'_{w',a} = 1$.*

*Proof.* We first show that any solution that meets these constraints is a solution to the LSR problem. The constraints about $T'$ being a refinement and copy-number consistency (constraint 1 in Problem 1) are explicitly enforced in this theorem. We must show that $T'$ is also a loss-supported phylogeny. The requirements for a loss-supported phylogeny is that every mutation occurs only once, which is enforced by (4) in this theorem, and that every mutation loss is supported. By (2), there are no mutation losses between cells that have the same copy-number state, and by (3) between cells with different copy-number states, mutation losses that are not supported are not allowed. Thus, $(T', B', \mathbf{c}')$ is a solution to the LSR.

We next show that any solution to the LSR also meets these constraints. We will do this by showing that any ($T'$, $B'$, $\mathbf{c}'$) that violates any one of these constraints cannot be a solution to the LSR.

1. $T'$ being a refinement and copy-number consistency (item (1)) are explicitly required by the LSR problem statement, and thus cannot be violated by any solution.
2. If $T'[\gamma(v)]$ is not a perfect phylogeny with respect to $B'[\gamma(v)]$, either there exists a mutation that occurs more than once in subtree $T'[\gamma(v)]$, in which case $T'$ is not a Dollo phylogeny, or there exists a mutation that is lost in subtree $T'[\gamma(v)]$. As every vertex in $T'[\gamma(v)]$ has the same copy-number state (by item (1)), this mutation loss is not supported. In either case it would mean $T'$ is not a loss-supported phylogeny and thus not a solution to the LSR problem.
3. If this condition is violated, then there is a mutation loss that is not supported. Thus, $T'$ is not a loss-supported phylogeny, and consequently not a solution to the LSR problem.
4. This condition must be true for $T'$ to be a Dollo, and thus a loss-supported phylogeny, and thus cannot be violated by any solution.

□

#### 3.2 Proof of Lemma 1

**Lemma 1.** *If there exists a solution to the LSR problem for a given $T$, $\mathbf{c}$, $\mathcal{L}$, $\mathbf{B}$, then there exists a solution $T'$, $\mathbf{c}'$, $\mathbf{B}'$ that meets the following conditions.*

*(1) For all $(v, w) \in E(T)$, $p(r(w))$ is a leaf of subtree $T'[\gamma(v)]$.*
*(2) For all $v \in V(T)$ such that $v$ is not the root of $T$, $b_{r(v)} = 1$ if $b_{v'} = 1$ for all $v' \in L(T'[\gamma(v)])$.*
*(3) For all $v \in V(T)$ and all loci $a$, $b_{p(r(v)),a} \geq b_{r(v),a}$.*

*Proof.* We will show by construction that for any solution $T'$, $\mathbf{c}'$, $\mathbf{B}'$ that violates these constraints, there exists another solution $T''$, $\mathbf{c}''$, $\mathbf{B}''$ that meets these constraints.

**(1) For all $(v, w) \in E(T)$, $p(r(w))$ is a leaf of subtree $T'[\gamma(v)]$.** Consider an edge $(p(r(w)), r(w)) \in E(T')$ such that $p(r(w))$ is not a leaf of $T'[\gamma(v)]$. That is, $p(r(w))$ has another child in $T'[\gamma(v)]$. We construct $T''$ by splitting $p(r(w))$ into two vertices $u$ and $u'$ such that there is an edge $(u, u') \in E(T'')$, $\mathbf{b}''_u = \mathbf{b}''_{u'}$ and $c''_u = c''_{u'}$, and the only outgoing edge from $u'$ is $(u', r(w))$. Thus, $u'$ is now the parent of $r(w)$ and $u'$ is a leaf. This split preserves the rest of the tree and does not introduce violations of any of the conditions in Theorem 1 or any of the other assumptions in this Lemma. Thus $T''$, $\mathbf{B}''$, $\mathbf{c}''$ is a solution to the LSR problem.

**(2) For all $v \in V(T)$ such that $v$ is not the root of $T$, $b_{r(v)} = 1$ if $b_{v'} = 1$ for all $v' \in L(T'[\gamma(v)])$.** Assume that $T', \mathbf{c}', \mathbf{B}'$ meet condition 1. If this constraint is violated, this means that there is some mutation $a$ that is gained in a subtree $T'[\gamma(v)]$ but there are no leaves of $T'[\gamma(v)]$ that do not contain $a$. Let $T'' = T'$, $c'' = c'$. Let $b'_{v',a} = 1$ if $v \in T'[\gamma(v)]$. This change does not violate any of the conditions in Theorem 1. Specifically, this change does not introduce new mutation gains, and as this only alters the mutation profiles of internal vertices of $T'[\gamma(v)]$ so this cannot introduce new mutation losses. As $T''$ and $\mathbf{c}''$ are preserved, refinement and copy-number consistency conditions are automatically met. This change may introduce violations to Assumption 3 in this Lemma that can subsequently be corrected as below.

**(3) For all $v \in V(T)$ and all loci $a$, $b_{p(r(v)),a} \geq b_{r(v),a}$.** This constraint states that there are no mutation gains on edges between subtrees. We construct $T''$ by performing a similar split as we did for constraint (1). Suppose there's an edge $(p(r(v)), r(v)) \in E(T')$ such that $b_{p(r(v)),a} = 0$ and $b_{r(v),a} = 1$. Split $p(r(v))$ into vertices $u, u'$ such that $b_{u,a} = 0$ and $b_{u',a} = 1$, and for all $a' \neq a$, $b_{u,a'} = b_{u',a'}$ and the only outgoing edge from $u'$ is $(u', r(w))$. This split preserves the rest of the tree and does not introduce violations of any of the conditions in Theorem 1 or any of the other assumptions in this Lemma. Thus $T''$, $\mathbf{B}''$, $\mathbf{c}''$ is a solution to the LSR problem. □

##### 3.3 Hardness of ML-LSR

**Lemma 2.** *The ML-LSR is NP-hard.*

*Proof.* We show this by reduction from the Flip problem which is known to be NP-Complete Chen et al. [2006].

**Problem 1.** *Given a binary matrix $\mathbf{B} \in \{0,1\}^{m \times n}$ and integer $\kappa \in \mathbb{N}$, decide whether there exists a directed perfect phylogeny matrix $\mathbf{B}' \in \{0,1\}^{m \times n}$ such that no more than $\kappa$ entries in $\mathbf{B}'$ differ from $\mathbf{B}$.*

Let $(\mathbf{B}, \kappa)$ be an instance of the Flip problem. For the corresponding instance of the ML-LSR problem, we let $k = 1$ and define the inputs as follows:

1. $T$ is the star phylogeny, where all leaves $v \in V(T)$ are attached to a single internal vertex;
2. $\mathbf{c} = \vec{1}$;
3. No mutation losses are supported in $\mathcal{L}$;
4. $\mathbf{X} = \mathbf{B}$, and $\mathbf{Y} = [1]^{m \times n}$.

Define a likelihood function $\Pr(X \mid Y, B^*)$ that is symmetric when $x_{v,a} \in \{0,1\}$ and $y_{v,a} = 1$

$$\log(\Pr(x_{v,a} \mid y_{v,a}, b_{v,a})) = \begin{cases} \alpha & \text{if } x_{v,a} = b_{v,a} \\ \beta & \text{if } x_{v,a} \neq b_{v,a} \end{cases} \tag{6}$$

such that $\beta < \alpha$. Thus the log-likelihood of a matrix $B*$ is

$$\log \Pr(X \mid Y, B^*) = \sum_{v=1}^{n} \sum_{a=1}^{m} \log(\Pr(x_{v,a} \mid y_{v,a}, b_{v,a})) \tag{7}$$

$$= \lambda \cdot \beta + (mn - \lambda)\alpha \tag{8}$$

$$\tag{9}$$

We claim that there exists a perfect phylogeny matrix $\mathbf{B}'$ with at most $\kappa$ changes if and only if there exists a solution $\mathbf{B}^*$ to the ML-LSR

$$\log \Pr(X \mid Y, B^*) \geq \kappa \cdot \beta + (mn - \kappa) \cdot \alpha \tag{10}$$

$$\tag{11}$$

We first show the forward direction. If there exists a perfect phylogeny matrix $\mathbf{B}'$ with at most $\kappa$ changes from $\mathbf{B}' = \mathbf{X}$, then the log-likelihood $\Pr(\mathbf{X} \mid \mathbf{Y}, \mathbf{B}') \geq \kappa \cdot \beta + (mn - \kappa) \cdot \alpha$. Thus for the maximum likelihood solution, $\mathbf{B}^*$, $\Pr(\mathbf{X} \mid \mathbf{Y}, \mathbf{B}^*) \geq \Pr(\mathbf{X} \mid \mathbf{Y}, \mathbf{B}')$.

We next show the reverse direction. If $\Pr(\mathbf{X} \mid \mathbf{Y}, \mathbf{B}^*) \geq \kappa \cdot \beta + (mn - \kappa)$ then $B^*$ has at most $\kappa$ changes from $\mathbf{X} = \mathbf{B}$. Thus, there exists a $\mathbf{B}' = \mathbf{B}^*$.

□

### 4 References

Edith M Ross and Florian Markowetz. Onconem: inferring tumor evolution from single-cell sequencing data. *Genome biology*, 17(1):69, 2016.

Katharina Jahn, Jack Kuipers, and Niko Beerenwinkel. Tree inference for single-cell data. *Genome biology*, 17(1):86, 2016.

Salem Malikic, Katharina Jahn, Jack Kuipers, S Cenk Sahinalp, and Niko Beerenwinkel. Integrative inference of subclonal tumour evolution from single-cell and bulk sequencing data. *Nature communications*, 10(1):2750, 2019a.

Jochen Singer, Jack Kuipers, Katharina Jahn, and Niko Beerenwinkel. Single-cell mutation identification via phylogenetic inference. *Nature communications*, 9(1):5144, 2018.

Simone Ciccolella, Mauricio Soto Gomez, Murray Patterson, Gianluca Della Vedova, Iman Hajirasouliha, and Paola Bonizzoni. Inferring cancer progression from single cell sequencing while allowing loss of mutations. *bioRxiv*, page 268243, 2018.

Mohammed El-Kebir. Sphyr: tumor phylogeny estimation from single-cell sequencing data under loss and error. *Bioinformatics*, 34(17):i671–i679, 2018.

Hamim Zafar, Anthony Tzen, Nicholas Navin, Ken Chen, and Luay Nakhleh. Sifit: inferring tumor trees from single-cell sequencing data under finite-sites models. *Genome biology*, 18(1):178, 2017.

Hamim Zafar, Nicholas Navin, Ken Chen, and Luay Nakhleh. Siclonefit: Bayesian inference of population structure, genotype, and phylogeny of tumor clones from single-cell genome sequencing data. *Genome Research*, 2019.

Salem Malikic, Farid Rashidi Mehrabadi, Simone Ciccolella, Md Khaledur Rahman, Camir Ricketts, Ehsan Haghshenas, Daniel Seidman, Faraz Hach, Iman Hajirasouliha, and S Cenk Sahinalp. Phiscs: a combinatorial approach for subperfect tumor phylogeny reconstruction via integrative use of single-cell and bulk sequencing data. *Genome Research*, 2019b.

Marco L Leung, Alexander Davis, Ruli Gao, Anna Casasent, Yong Wang, Emi Sei, Eduardo Sanchez, Dipen Maru, Scott Kopetz, and Nicholas E Navin. Single cell dna sequencing reveals a late-dissemination model in metastatic colorectal cancer. *Genome research*, pages gr–209973, 2017.

Duhong Chen, Oliver Eulenstein, David Fernandez-Baca, and Michael Sanderson. Minimum-flip supertrees: complexity and algorithms. *IEEE/ACM Transactions on Computational Biology and Bioinformatics (TCBB)*, 3(2):165–173, 2006.
